# The evolutionary basis of premature migration in Pacific salmon highlights the utility of genomics for informing conservation

**DOI:** 10.1101/056853

**Authors:** Daniel J. Prince, Sean M. O’Rourke, Tasha Q. Thompson, Omar A. Ali, Hannah S. Lyman, Ismail K. Saglam, Thomas J. Hotaling, Adrian P. Spidle, Michael R. Miller

**Author notes:** These authors contributed equally to this work (Sean M. O’Rourke, Tasha Q. Thompson). **Corresponding Author:** Michael R. Miller, Department of Animal Science, University of California, 1 Shields Ave, Davis CA 95616 USA.

## Abstract

The delineation of conservation units (CUs) is a challenging issue that has profound implications for minimizing the loss of biodiversity and ecosystem services. CU delineation typically seeks to prioritize evolutionary significance and genetic methods play a pivotal role in the delineation process by quantifying overall differentiation between populations. While CUs that primarily reflect overall genetic differentiation do protect adaptive differences between distant populations, they do not necessarily protect adaptive variation within highly connected populations. Advances in genomic methodology facilitate the characterization of adaptive genetic variation, but the potential utility of this information for CU delineation is unclear. Here we use genomic methods to investigate the evolutionary basis of premature migration in Pacific salmon, a complex behavioral and physiological adaptation that exists within highly-connected populations and has experienced severe declines. Strikingly, we find that premature migration is associated with the same single locus across multiple populations in each of two different species. Patterns of variation at this locus suggest that the premature migration alleles arose from a single evolutionary event within each species and were subsequently spread to distant populations through straying and positive selection. Our results reveal that complex adaptive variation can depend on rare mutational events at a single locus, demonstrate that CUs reflecting overall genetic differentiation can fail to protect evolutionarily significant variation that has substantial ecological and societal benefits, and suggest that a supplemental framework for protecting specific adaptive variation will sometimes be necessary to prevent the loss of significant biodiversity and ecosystem services.

## Introduction

Invaluable economic, ecological, and cultural benefits are being lost worldwide as biodiversity decreases due to human actions (Millennium Ecosystem Assessment 2005; Krauss *et al.* 2010; Pimm *et al.* 2014). Legislation exists in many countries throughout the world that provides a framework to protect unique species as well as population segments below the species level (Allendorf et al. 2012). Protection is achieved by assessing the health of a defined conservation unit and, if the unit is at risk, critical habitat is preserved/restored and stressors are restricted until the risk is eliminated. Assessing risk and developing a protection strategy is not possible without first establishing unit boundaries. Since the number of units that can be effectively managed is resource-limited (Waples 1998), the delineation of units must be strategic and prioritize evolutionary significance (Ryder 1986; Waples 1991; Dizon *et al.* 1992; Moritz 1994; Crandall *et al.* 2000; Fraser and Bernatchez 2001). Several criteria, such as genetic and ecological exchangeability (Crandall *et al.* 2000), have been proposed for assessing evolutionary significance for conservation unit delineation, but directly evaluating these criteria in natural populations is difficult (Allendorf *et al.* 2012).

Genetic methods play a pivotal role in evaluating evolutionary significance for conservation unit delineation (Crandall *et al.* 2000). To this end, genetic data from different regions of the genome are combined to produce measurements of overall genetic differentiation between populations. These measurements represent typical regions of the genome and are used as a proxy for evolutionary significance in conservation unit delineation. However, because most genomic regions are primarily influenced by gene flow and genetic drift as opposed to selection, these measurements may fail to account for important adaptive differences between populations (Funk *et al.* 2012). Recent advances in genetic methodology facilitate the identification and evolutionary analysis of adaptively important loci (Colosimo *et al.* 2005; Miller *et al.* 2007; Baird *et al.* 2008; Hohenlohe *et al.* 2010; Miller *et al.* 2012; Pearse *et al.* 2014; Barson *et al.* 2015; Ali *et al.* 2016) and provide an alternative way to assess evolutionary significance, but the utility of these loci for conservation unit delineation is unclear and disputed (Allendorf *et al.* 2010; Funk *et al.* 2012; Shafer *et al.* 2015, 2016; Garner *et al.* 2016; Pearse 2016).

Pacific salmon (*Oncorhynchus* spp.) provide a unique opportunity to investigate the application of genetic tools to the conservation of biodiversity below the species level (Waples 1991, 1998; Allendorf *et al.* 1997; Pennock and Dimmick 1997; Wainwright and Waples 1998). Despite extensive conservation efforts, Pacific salmon have been extirpated from almost forty percent of their historical range in the contiguous U.S. while many remaining populations have experienced dramatic declines and face increasing challenges from climate change (Levin and Schiewe 2001; Augerot 2005; Gustafson *et al.* 2007; Moyle *et al.* 2008; Muñoz *et al.* 2015). Reintroduction attempts of extirpated populations are largely unsuccessful as precise natal homing across highly heterogeneous environments has resulted in divergent selection and abundant local adaptation (Taylor 1991; Fraser *et al.* 2006; Miller *et al.* 2012; Anderson *et al.* 2014). Thus, maintaining existing stocks is critical for preserving the species themselves as well as the communities and ecosystems which rely on their presence (Committee on Protection and Management of Pacific Northwest Anadromous Salmonids *et al.* 1996). Genetic methods have been used extensively in the process of delineating conservation units in Pacific salmon (called evolutionarily significant units [ESUs] or distinct population segments [DPSs] depending on the species) and, as a consequence of patterns of gene flow, have resulted in units that primarily reflect geography (Busby *et al.* 1996; USFWS and NOAA 1996; Myers *et al.* 1998; NOAA 2006). While current ESUs and DPSs certainly protect adaptive differences between distant populations, adaptations within highly connected populations are not necessarily protected (Crandall *et al.* 2000; Moyle *et al.* 2008). However, the evolutionary significance of these adaptations and the potential long-term consequences of not independently protecting them are poorly understood.

Perhaps the most recognized example of differential adaptation within highly connected populations of Pacific salmon is variation in adult migration timing (also called run timing) (Moyle 2002; Quinn 2005; Quinn *et al.* 2015). In contrast to typical adult salmon that mature sexually before freshwater migration, premature migrating individuals have a complex behavioral and physiological adaptation which allows them to access distinct habitats and spawn earlier in the season (Quinn *et al.* 2015). Premature migrating populations also distribute ocean-derived nutrients higher into watersheds and provide additional, more-coveted, and culturally important harvest opportunities due to their distinct migration time and high fat content (McEvoy 1990; Hearsey and Kinziger 2014). Due to their extended time in freshwater and reliance on headwater habitat, premature migrating populations have suffered grossly disproportionate impacts from human actions such as dam building, mining, and logging (Busby *et al.* 1996; Myers *et al.* 1998; Kareiva *et al.* 2000; Committee on Endangered and Threatened Fishes in the Klamath River Basin *et al.* 2004; Moyle *et al.* 2008; Williams *et al.* 2013; Quinn *et al.* 2015). Genetic analyses typically find little differentiation between proximate premature and mature migrating populations (Allendorf 1975; Chilcote *et al.* 1980; Thorgaard 1983; Nielsen and Fountain 1999; Waples *et al.* 2004; Clemento III 2006; Kinziger *et al.* 2013; Brieuc *et al.* 2015; Arciniega *et al.* 2016), and as a result, they are generally grouped into the same ESU or DPS (Busby *et al.* 1996; Myers *et al.* 1998). Therefore, despite the extirpation or substantial decline of premature migrating populations, the ESUs or DPSs to which they belong usually retain relatively healthy mature migrating populations and thus have low extinction risk overall (Busby *et al.* 1996; Myers *et al.* 1998; Williams *et al.* 2013). Here we investigate the genetic and evolutionary basis of premature migration to explore potential consequences of not independently protecting this beneficial adaptation as well as the utility of genomics for informing conservation.

## Results

### Initial genomic analysis consistent with current steelhead DPS delineations

Dramatic examples of premature migration are observed in coastal (non-interior) populations of steelhead (anadromous rainbow trout; *O. mykiss*) and Chinook salmon (*O. tshawytscha*). In these populations, premature migrating individuals (called summer steelhead or spring Chinook) utilize receding spring flows during freshwater migration to reach upstream habitat prior to hostile summer conditions in the lower watershed, hold for several months in deep cool pools while their gametes mature, then spawn at similar times to mature migrating individuals which have just entered freshwater (Moyle 2002; Quinn *et al.* 2015). We began our investigation by compiling a set of 148 steelhead samples from five coastal locations across four DPSs in California and Oregon (Figure 1A). Four of the locations (Eel, New, Siletz, and North Umpqua) represent the few remaining watersheds with significant wild premature migrating populations. The fifth location, Scott, contains only mature migrating individuals. Our sampling focused as much as possible on individuals that could be confidently categorized as premature or mature migrating based on collection date and location (Figure 1B; Table S1).

**Figure 1.**
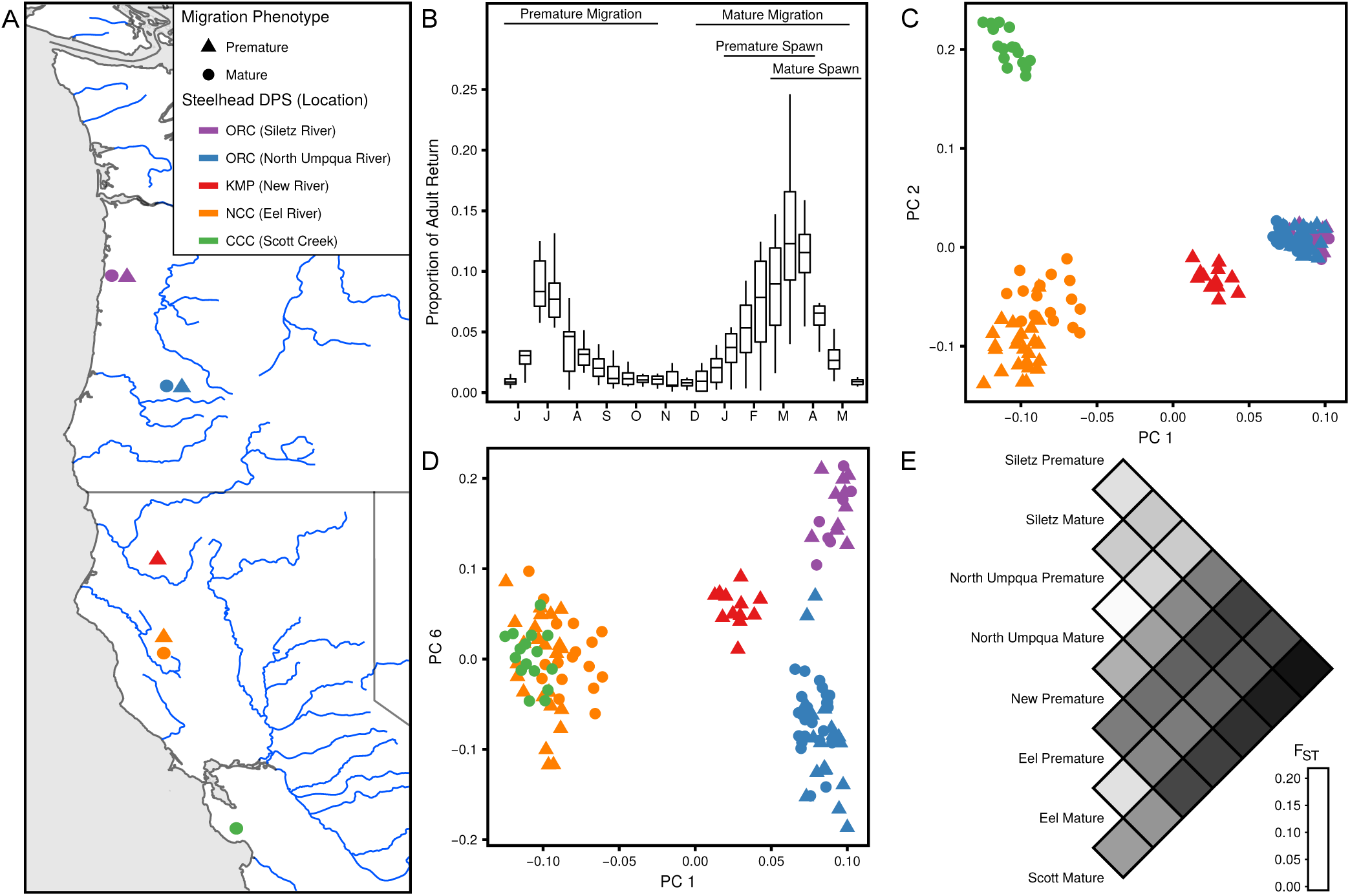
Genetic structure of premature and mature migrating steelhead populations. A) Map of steelhead sample locations and migration phenotypes; color indicates location, shape indicates migration phenotype. B) Bimonthly proportion of annual adult steelhead return over Winchester Dam on the North Umpqua River (2003 - 2013); horizontal bars depict migration and spawn timing of premature and mature migrating populations. C) and D) Principal component analysis and E) pairwise FST estimates using genome-wide SNP data.

To collect high-resolution genomic information from these samples, we prepared individually barcoded restriction-site associated DNA (RAD) libraries, sequenced them using paired-end Illumina technology, and aligned the sequence reads to a recent draft of the rainbow trout genome (Berthelot *et al.* 2014; Table S2, S1). We then used a probabilistic framework to discover SNPs and genotype them in each individual (Korneliussen *et al.* 2014). A total of 9,864,960 genomic positions were interrogated in at least 50% of individuals and 615,958 SNPs (i.e. segregating sites) were identified (p-value < 10^-6^). Of these SNPs, 215,345 had one genotype posterior greater than 0.8 in at least 50% of individuals. Population structure characterization and genome-wide analyses in nonmodel organisms are typically carried out with far fewer SNPs (Narum *et al.* 2013). We conclude that the sequence data obtained is appropriate for genome-wide measurements as well as high-resolution analyses of specific genomic regions.

To characterize the genetic structure of these populations, we performed a principal component analysis and estimated pairwise FST using genome-wide genotype data (Fumagalli *et al.* 2014). The first two principal components (PCs) revealed four distinct groups corresponding to the four current DPSs (Figure 1C). The Siletz and North Umpqua, which are two distinct locations within the Oregon Coast DPS, did not break into distinct groups until PC6 (Figure 1D), indicating relatively low genetic differentiation between distinct locations within a DPS. In all cases, individuals with different migration phenotypes from the same location were in the same group. The pairwise FST estimates also revealed strong genetic differentiation between locations but little differentiation between migration phenotypes from the same location (Figure 1E). The mean pairwise FST between migration groups from the same location was 0.032 (range: 0.018 - 0.039, n = 3) whereas the mean between groups from different locations was 0.125 (range: 0.049 - 0.205, n = 25). The combination of this genetic structure and observations of hybridization between premature and mature migrating individuals (Chilcote *et al.* 1980) suggests higher rates of gene flow between different migration groups from the same location than between groups from different locations. Thus, as found in previous analyses, the overall genetic structure among steelhead populations is predominantly influenced by geography as opposed to migration phenotype. We conclude that measurements of overall genetic differentiation from genome-wide SNP data are consistent with current steelhead DPS delineations.

### Premature migrating steelhead explained by a single allelic evolutionary event at a single locus

To identify genomic loci associated with premature migration, we performed association mapping of migration category. We used a likelihood ratio test (Kim *et al.* 2011) with lambda correction for population stratification (Price *et al.* 2010) to compare 181,954 SNPs between migration categories in the North Umpqua and found 14 SNPs that were significant (p-value < Bonferroni-corrected α of 0.05). Strikingly, all of these SNPs were located within a 211,251 bp region (568,978 - 780,229) on a single 1.95 Mbp scaffold (Figure 2A, S1A, S1B; Table S3). Furthermore, when this analysis was repeated with Eel individuals using 170,678 SNPs, we obtained a similar pattern of association (Figure 2B, S1C, S1D; Table S3). The strongest associated SNPs in both sample locations were flanking two restriction sites approximately 50 kb apart and located just upstream and within a gene identified as *GREB1L* (Figure 2C; see Discussion for more information on *GREB1L*). The strength of these associations was unexpected given the phenotypic complexity of premature migration and the relatively low number of samples analyzed. We conclude that the same single locus is strongly associated with migration phenotype in at least two DPSs.

**Figure 2.**
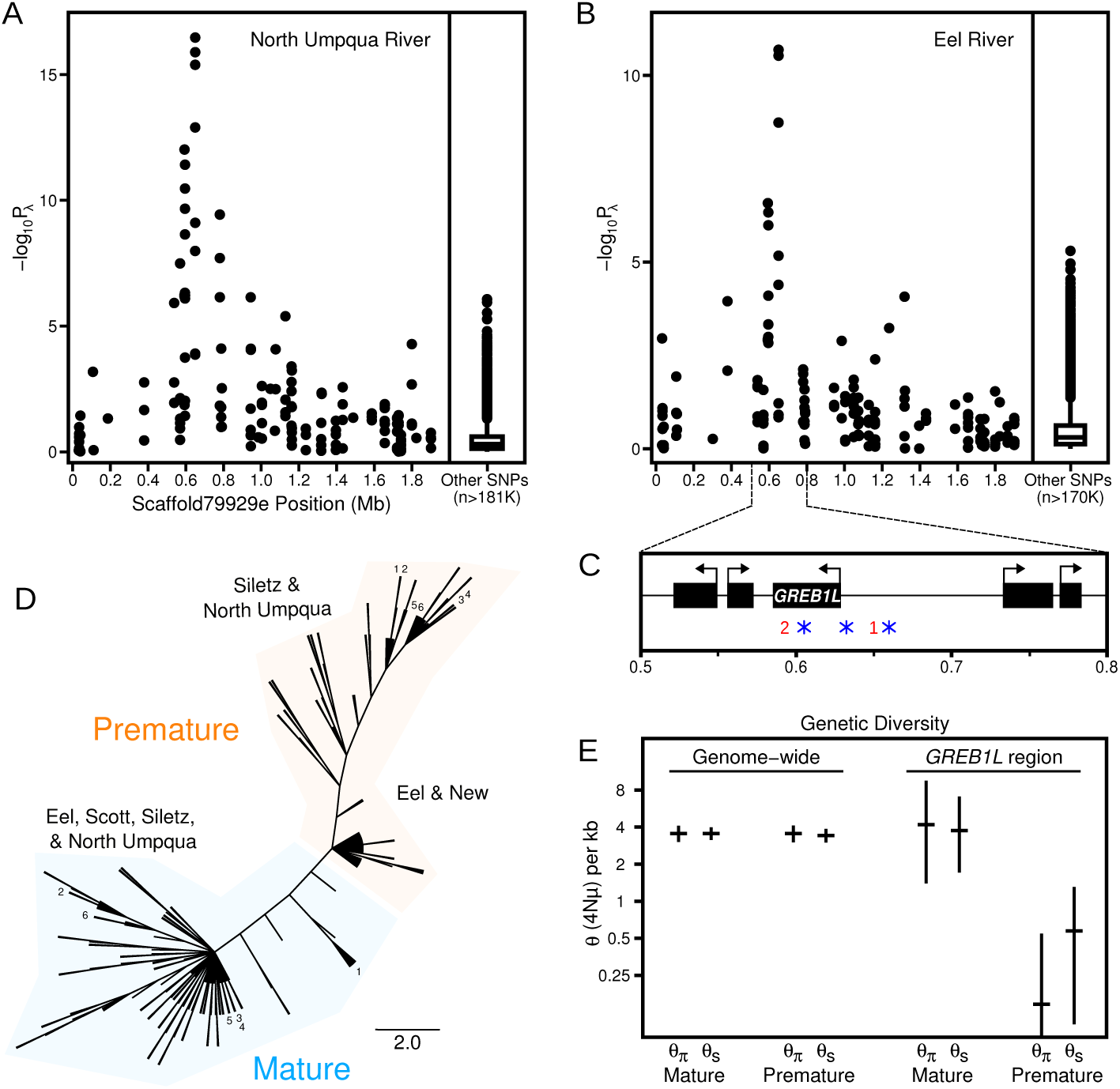
Genetic and evolutionary basis of premature migration in steelhead. Association mapping of migration category in A) North Umpqua River and B) Eel River steelhead. C) Gene annotation of region with strong association; red numbers indicate genomic positions of two restriction sites flanked by strongest associated SNPs, blue asterisks indicate positions of amplicon sequencing. D) Phylogenetic tree depicting maximum parsimony of phased amplicon sequences from all individuals; branch lengths, with the exception of terminal tips, reflect nucleotide differences between haplotypes; numbers identify individuals with one haplotype in each migration category clade. E) Genome-wide and *GREB1L* region diversity estimates in North Umpqua for each migration category with 95% confidence intervals from coalescent simulations.

To investigate the evolutionary history of this locus, we sequenced three amplicons, each of approximately 500 bp, from the *GREB1L* region in all individuals (Figure 2C; Table S4, S5, S1) and used these sequences to construct a haplotype tree based on parsimony (Felsenstein 1989). Strikingly, the tree contained two distinct monophyletic groups corresponding to migration phenotype (Figure 2D). For 123 out of 129 individuals, both haplotypes separated into the appropriate migration category clade. The remaining six individuals (4 Siletz and 2 North Umpqua samples originally classified as mature migrating) had one haplotype in each migration category clade (Figure 2D), suggesting heterozygosity at the causative polymorphism(s). Furthermore, although there was little differentiation within the mature migration clade, premature migration haplotypes from the Siletz and North Umpqua were more divergent from the mature migration clade than those from the Eel and New (Figure 2D; see Discussion for more information on heterozygotes and differentiation within the premature clade). The overall tree topology is inconsistent with premature migration alleles originating from independent evolutionary events in different locations. We conclude that there is a complete association between variation at this locus and migration category and that the premature migration alleles from all locations arose from a single evolutionary event.

To examine the evolutionary mechanisms leading to the dispersal of the premature migration allele as well as reconcile the difference between patterns of variation at the *GREB1L* locus and overall genetic structure, we summarized patterns of genetic variation using two estimators of theta (θ = 4Nμ). One estimator is based on average pairwise differences (θπ; Tajima 1983) and the other is based on number of segregating sites (θ_s_; Watterson 1975). When genome-wide data were used, both estimators produced similar thetas for each migration category (Figure 2E). The *GREB1L* region of mature migrating individuals also produced thetas similar to the genome-wide analysis. However, premature migrating individuals from the North Umpqua had strikingly lower thetas (Figure 2E) and a significantly skewed site frequency spectrum (Tajima’s D = −2.08, p-value = 0.001; Tajima 1989) indicative of strong, recent positive selection in the *GREB1L* region. Premature migrating individuals from the Eel also had reduced thetas in the *GREB1L* region (premature: θ_π_/kb = 2.48, θ_S_/kb = 2.67; mature: θ_π_/kb = 3.59, θ_S_/kb = 4.00), but the site frequency spectrum was not significantly skewed, consistent with an older selection event. While both demography and selection can reduce nucleotide diversity and skew the site frequency spectrum, this pattern is specific to the *GREB1L* region as opposed to genome-wide, implicating selection as the cause. Furthermore, the combination of a stronger signature of selection and a more divergent sequence pattern in the northern premature migration haplotypes is consistent with a relatively recent (perhaps since the last glacial maximum) movement of the premature migration allele to the north. We conclude that, upon entering new locations via straying, positive selection allowed the premature migration allele to persist despite reductions in overall genetic differentiation due to hybridization with local mature migrating populations.

### Premature migrating Chinook also explained by a single allelic evolutionary event in GREB1L region

To broaden our investigation into premature migration, we compiled a set of 250 Chinook samples from nine locations across five ESUs in California, Oregon, and Washington (Figure 3A). Similar to steelhead, our sampling focused as much as possible on individuals that could be confidently categorized as premature or mature migrating based on collection time and location (Table S6). We then prepared individually barcoded RAD libraries, sequenced them using paired-end Illumina technology, and aligned the sequence reads to the same rainbow trout reference assembly used above (Table S7, S6). No reference genome is available for Chinook, and rainbow trout, which diverged from Chinook approximately 10-15 mya (Crête-Lafrenière *et al.* 2012; Zhivotovsky 2015), is the closest relative with a draft genome assembly. Using the methods described above, a total of 3,910,009 genomic positions were interrogated in at least 50% of individuals and 301,562 SNPs were identified (p-value < 10^-6^). Of these SNPs, 55,797 had one genotype posterior greater than 0.8 in at least 50% of individuals. Although the alignment success was lower and subsequent SNP discovery and genotyping produced fewer SNPs compared to steelhead, the large number of SNPs discovered and genotyped should still be adequate for downstream analysis.

To characterize the genetic structure of these populations, we performed a principal component analysis and estimated pairwise FST using the genotype information described above. The first two PCs revealed four groups: the largest group contained all coastal ESUs, the second contained the two Puget Sound ESU locations, and the last two groups corresponded to the two locations within the Upper Klamath-Trinity Rivers ESU and were only differentiated by the second axis (Figure 3B). In all cases, individuals from the same location but with different migration phenotypes were in the same group, and locations within groups became differentiated as additional PCs were examined (data not shown). The mean pairwise FST between migration categories from the same location was 0.037 (range: 0.009 – 0.093, n = 7) and the mean between groups from different locations was 0.097 (range: 0.021 – 0.199, n = 113; Figure 3C). Thus, similar to what we found in steelhead, the overall genetic structure is strongly influenced by geography as opposed to migration phenotype. We conclude that measurements of overall genetic differentiation from genome-wide SNP data are consistent with current Chinook ESUs.

**Figure 3.**
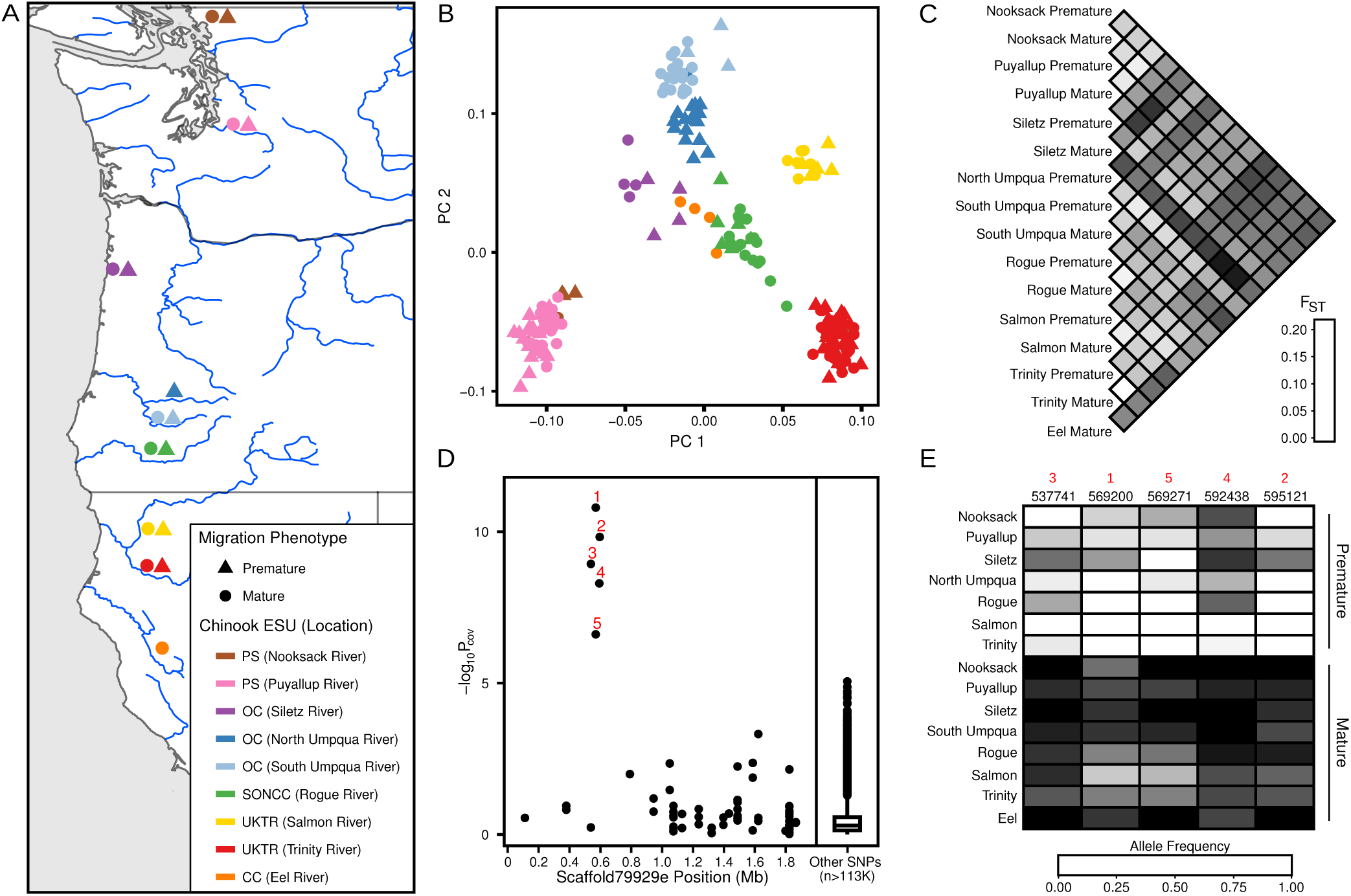
Genetic and evolutionary basis of premature migration in Chinook. A) Map of Chinook sample locations and migration phenotypes; color indicates location, shape indicates migration category. B) Principal component analysis and C) pairwise FST estimates using genome-wide SNP data. D) Association mapping of migration category in Chinook; red numbers indicate significant SNPs. E) Allele frequency shift at significant SNPs between premature and mature migrating populations. Black numbers indicate SNP position on scaffold79929e.

To investigate the genetic architecture and evolutionary basis of premature migration in Chinook, we conducted association mapping with 114,036 SNPs using a generalized linear framework with covariate correction for population stratification (Price *et al.* 2010; Skotte *et al.* 2012). Strikingly, we again found a single significant peak of association (p-value < Bonferroni-corrected α of 0.05) that contained five SNPs within 57,380 bp (537,741 – 595,121) in the same *GREB1L* region identified in steelhead (Figure 3D; Table S8). We next examined allele frequencies at these five SNPs and found a strong and consistent shift between all premature and mature migrating populations independent of location (Figure 3E). Thus, despite having lower genomic resolution and fewer samples per location, these results demonstrate that the *GREB1L* region is also the primary locus associated with premature migration in Chinook. Furthermore, the shift of allele frequencies in the same direction across all populations is inconsistent with the premature migration alleles in Chinook being a product of multiple independent evolutionary events. Although the genomic region was consistent between species, the SNPs identified in Chinook were distinct from those in steelhead (Table S8, S3). In other words, the premature and mature migrating Chinook haplotypes are more similar to each other than to either of the steelhead haplotypes and vice versa, suggesting independent allelic evolutionary events in each species. We conclude that the same evolutionary mechanism used in steelhead, with a single allelic evolutionary event in the *GREB1L* region that subsequently spread to different locations, also explains premature migration in Chinook.

## Discussion

Our association analysis across multiple populations in each of two different species as well as an independent analysis on Klickitat River steelhead (Hess *et al.* 2016) suggests that either the function or regulation of *GREB1L* is modified in premature migrating individuals. *GREB1L* is highly conserved across vertebrates, encodes a nuclear hormone receptor coactivator, and is differentially regulated by feeding and fasting in AgRP neurons of the hypothalamic arcuate nucleus in mice (Mohammed *et al.* 2013; Henry *et al.* 2015). The strength of the associations as well as the known role of AgRP neurons in modulating diverse behavior and metabolic processes such as foraging and fat storage (Dietrich *et al.* 2015; Henry *et al.* 2015) provides evidence for and an explanation as to how the complex premature migration phenotype could be controlled by this single locus. An alternative explanation is that the *GREB1L* region only influences a subset of the phenotypic components of premature migration and that other important loci were not identified due to technical or biological reasons. Regardless, our results indicate that an appropriate genotype at this locus is necessary for successful premature migration.

Given that premature migration alleles at this locus are critical for premature migration, our results on the evolutionary history of these alleles provide important insights into the potential for premature migration to persist during declines and reemerge if lost. Finding that the same locus is associated with premature migration in both steelhead and Chinook indicates that genetic mechanisms capable of producing this phenotype are very limited. Although some loci can be predisposed to functionally-equivalent mutations in relatively short evolutionary time scales (Cockram *et al.* 2007; Chan *et al.* 2010), this does not appear to be the case with the *GREB1L* region. In predisposed loci, several independent mutations with the same phenotypic effect are observed in different populations of a single species (Cockram *et al.* 2007; Chan *et al.* 2010). In contrast, we observed only one evolutionary event that produced a premature migration allele in each species despite the 10-15 million years since they diverged (Crête-Lafrenière *et al.* 2012; Zhivotovsky 2015). This suggests the *GREB1L* region is not predisposed to mutational events that create new premature migration alleles and that such allelic evolutionary events are exceptionally rare. Thus, the premature migration allele and phenotype cannot be expected to readily re-evolve if the alleles are lost.

The rarity of mutational events that produce premature migration alleles at this locus highlights the importance of existing premature migration alleles. Unlike alleles with a small effect on phenotype, alleles with a large effect on phenotype are expected to be rapidly lost from a population when there is strong selection against the phenotype they promote (Charlesworth and Charlesworth 2010). An important exception to this is when an allele is recessive and therefore masked in the heterozygous state (Colosimo *et al.* 2005; Charlesworth and Charlesworth 2010). Thus, the dominance pattern of the *GREB1L* locus has critical implications for the persistence of premature migration alleles during declines of the premature migration phenotype. Although our sampling focused on migration peaks (Figure 1B) and was not designed to investigate the migration phenotype of heterozygotes, the recently published Klickitat data (Hess *et al.* 2016) included samples collected outside the migration peaks. Strikingly, a reanalysis of these data suggests that the same haplotype is associated with premature migration (Figure 4A; Table S3) and that heterozygotes display an intermediate phenotype (Figure 4B, S2). This explains the high frequency of heterozygotes in our Siletz mature migrating samples (4 out of 10) which were collected prior to the peak of mature migration and far upstream in the watershed (Table S1). Thus, the premature migration allele does not appear to be masked in the heterozygous state and cannot be expected to be maintained as standing variation in populations that lack the premature migration phenotype.

**Figure 4.**
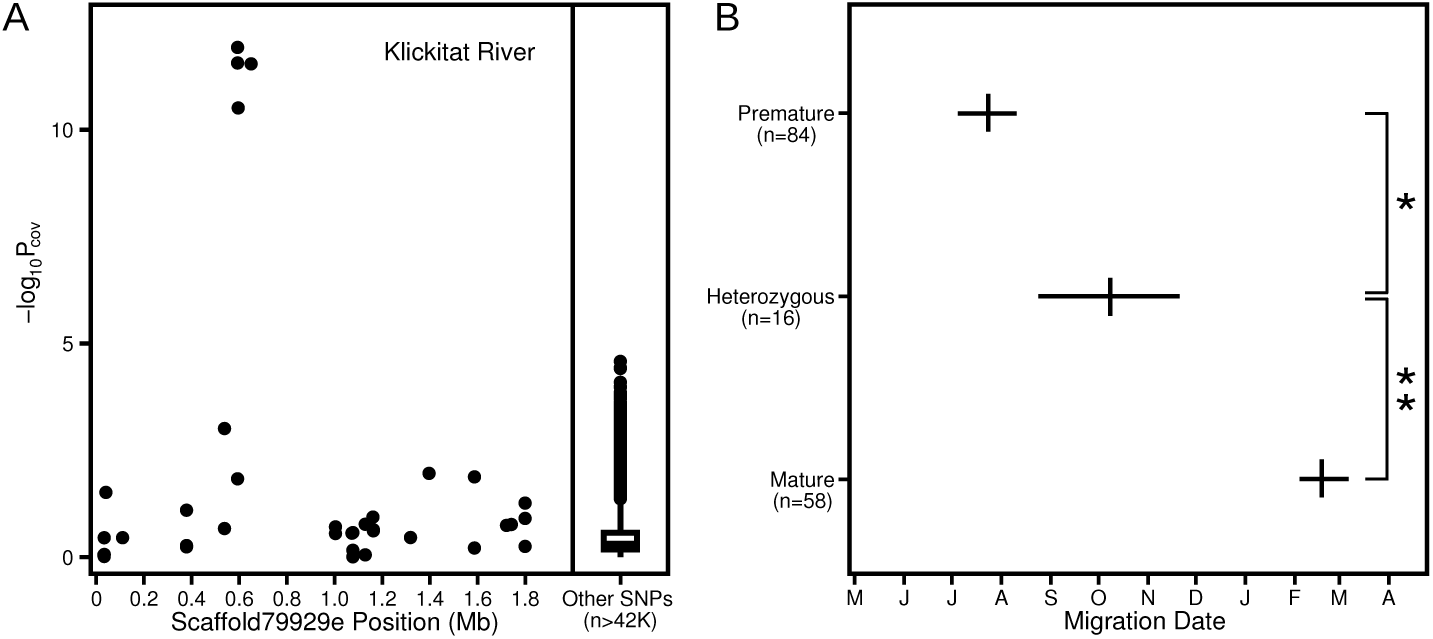
Dominance pattern of *GREB1L* locus. A) Association mapping of migration date in Klickitat River steelhead. B) Migration date mean and 95% confidence interval of individuals categorized as homozygous for the premature migration allele, heterozygous, and homozygous mature. * p-value = 0.00574; ** p-value = 2.95×10^-5^.

Two additional lines of evidence suggest that the premature migration allele will not be maintained as standing variation in mature migrating populations. First, the combination of the strong bimodal phenotypic distribution that is usually observed (e.g. Figure 1B) and the ecology of premature migration (see introduction; Moyle 2002; Quinn *et al.* 2015) suggests a general pattern of disruptive selection against individuals with an intermediate phenotype (e.g. heterozygotes). Although heterozygotes are expected to be produced by hybridization in locations where both migration categories exist (e.g. we observed two heterozygotes in the North Umpqua which has the lowest genetic differentiation between migration groups; Figure 1E), their presence does not suggest that the premature migration allele will be maintained by mature migrating populations. Second, the genetic differentiation between premature migration haplotypes from California and Oregon steelhead (Figure 2D) indicates that, unlike mature migration alleles, premature migration alleles are not freely moving across this area. This result reveals that mature migrating populations do not act as an influential source or conduit of premature migration alleles despite being abundant and broadly distributed. Therefore, premature migrating populations appear ultimately necessary for both the maintenance and spread of these alleles.

Previously, studies revealing that overall genetic structure among populations of steelhead and Chinook primarily reflects geography (as opposed to migration phenotype) suggested that premature migration evolved independently in many locations within each species (Thorgaard 1983; Waples *et al.* 2004; Arciniega *et al.* 2016). This implied that premature migration is evolutionarily replaceable over time frames relevant to conservation planning (e.g. tens to hundreds of years; Waples *et al.* 2004) and is not an important component in the evolutionary legacy of the species (Williams *et al.* 2013). Although these interpretations were logical given the data available at that time, our results demonstrate that the evolution was not independent in each location but instead relied on pre-existing genetic variation. Thus, although evolving the premature migration phenotype in new locations could be rapid if robust premature migrating populations are present in proximate locations, the widespread extirpation and decline of premature migrating populations (Busby *et al.* 1996; Myers *et al.* 1998; Kareiva *et al.* 2000; Committee on Endangered and Threatened Fishes in the Klamath River Basin *et al.* 2004; Moyle *et al.* 2008; Williams *et al.* 2013; Quinn *et al.* 2015) has greatly diminished the potential restoration and expansion (e.g. into new habitats that become available with climate change) of premature migration across at least a substantial proportion of the range for both species (Miller *et al.* 2012).

The combination of three key results from this study has broad conservation implications which highlight the utility of genomics for informing conservation. First, we present a striking example of how a single allele at a single locus can have significant economic, ecological, and cultural importance. Second, we show that mutations producing an important allele can be very rare from an evolutionary perspective, suggesting the allele will not readily re-evolve if lost. Lastly, we observe that patterns of significant adaptive allelic variation can be completely opposite from patterns of overall genetic differentiation. Taken together, our results demonstrate that conservation units reflecting overall genetic differentiation can fail to protect evolutionarily significant variation that has substantial ecological and societal benefits, and suggest that a supplemental framework for protecting specific adaptive variation will sometimes be necessary to prevent the loss of significant biodiversity and ecosystem services.

## Methods

### Sample collection and molecular biology

Fin clips were taken from live adults or post-spawn carcasses (Table S1, S6), dried on Whatman qualitative filter paper (Grade 1), and stored at room temperature. DNA was extracted with either the DNeasy Blood & Tissue Kit (Qiagen) or a magnetic bead-based protocol (Ali *et al.* 2016) and quantified using Quant-iT PicoGreen dsDNA Reagent (Thermo Fisher Scientific) with an FLx800 Fluorescence Reader (BioTek Instruments).

SbfI RAD libraries were prepared with well and plate (when applicable) barcodes using either the traditional or new RAD protocol (Ali *et al.* 2016) and sequenced with paired-end 100 bp reads on an Illumina Hiseq 2500 (Table S2, S7). In some cases, the same sample was included in multiple libraries to improve sequencing coverage.

For amplicon sequencing, genomic DNA extractions were rearrayed into 96-well plates and diluted 1:40 with low TE pH 8.0 (10 mM Tris-HCl, 0.1 mM EDTA). 2 μl of this diluted sample was used as PCR template for each of three amplicons in the *GREB1L* region (Figure 2; Table S4). Multiple forward primers were synthesized for each amplicon. Each forward primer contained a partial Illumina adapter sequence, a unique inline plate barcode, and the amplicon specific sequence (Table S4, S5). Initial PCR reactions were performed in 96-well plates using OneTaq DNA polymerase (New England Biolabs) at the recommended conditions with an annealing temperature of 61°C and 35 cycles. These reaction plates were then combined into a single plate that preserved the well locations. The pooled PCR products were cleaned with Ampure XP beads (Beckman Coulter) and a second round of PCR with 8 cycles was performed to add the remaining Illumina adapter sequence and a unique TruSeq barcode to each well (Table S4, S5). From each final PCR reaction, 2 μl was removed, pooled, and purified with Ampure XP beads. The final amplicon library was sequenced with paired-end 300 bp reads on an Illumina MiSeq.

### RAD analysis

RAD sequencing data were demultiplexed by requiring a perfect barcode and partial restriction site match (Ali *et al.* 2016). Sequences were aligned to a slightly modified version of a recent rainbow trout genome assembly (Berthelot *et al.* 2014; see scaffold79929e assembly and annotation) using the backtrack algorithm of BWA (Li and Durbin 2009) with default parameters. SAMtools (Li *et al.* 2009) was used to sort, filter for proper pairs, remove PCR duplicates, and index BAM files (Table S2, S7). In cases where the same sample was sequenced in multiple libraries, BAM files from the same sample were merged prior to indexing using SAMtools (Table S1, S2, S6, S7).

Additional BAM file sets were generated to account for technical variation among samples. To minimize variation associated with the two distinct library preparation protocols used in Chinook (Ali *et al.* 2016; Table S7), a set of single-end BAM files were generated for Chinook that contained only trimmed reads from the restriction-site end of the RAD fragments. To prepare these files, these reads were trimmed to 75 bp from the 3' end after removing 5 bp from the 5' end. Next, paired-end alignments were performed and processed as above. Lastly, reads from the variable end of RAD fragments were removed (Table S7). To remove variation associated with variable sequencing depth, a set of subsampled BAM files were generated by using SAMtools to randomly sample approximately 120,000 alignments from paired-end BAM files for steelhead and approximately 60,000 alignments from single-end BAM files for Chinook.

All RAD analyses were performed using ANGSD (Korneliussen *et al.* 2014) with a minimum mapping quality score (minMapQ) of 10, a minimum base quality score (minQ) of 20, and the SAMtools genotype likelihood model (GL 1; Li 2011). Unless otherwise noted, samples with less alignments than required for subsampling were excluded (Table S1, S6), and only sites represented in at least 50% of the included samples (minInd) were used.

Principal component analyses (PCA) and association mapping were performed by identifying polymorphic sites (SNP _pval 1e-6), inferring major and minor alleles (doMajorMinor 1; Skotte *et al.* 2012), estimating allele frequencies (doMaf 2; Kim *et al.* 2011), and retaining SNPs with a minor allele frequency of at least 0.05 (minMaf). For PCA, subsampled BAM files were used and genotype posterior probabilities were calculated with a uniform prior (doPost 2). The ngsCovar (Fumagalli *et al.* 2013) function implemented in ngsTools (Fumagalli *et al.* 2014) was used to calculate a covariance matrix from called genotypes. For association mapping, paired-end BAM files were used with two distinct tests. The frequency test with known major and minor alleles (doAsso 1) implements a likelihood ratio test using read counts (Kim *et al.* 2011). This test has good statistical power even with lower coverage data but does not allow the inclusion of covariates to correct for population stratification. The score test (doAsso 2) uses a generalized linear framework on posterior genotype probabilities (Skotte *et al.* 2012). This test allows the inclusion of covariates to correct for population stratification but has less statistical power than the frequency test. For the Umpqua and Eel steelhead associations, the frequency test with lambda correction for population stratification (Price *et al.* 2010) was used because there were relatively few samples and weak population structure. Lambda is the ratio of observed and expected median chi-squared values and used to correct the observed chi-squared values before converting them to p-values (Price *et al.* 2010; Figure S1A, S1C; Table S3). For the Chinook association, the score test with covariate correction for population stratification was used because there were many samples and complex population structure. The positions of each sample along the first 15 principal components (PCs) were used as covariates.

Genome-wide F_ST_ between population pairs was estimated by first estimating a site frequency spectrum (SFS) for each population (doSaf; Nielsen *et al.* 2012) using paired-end BAM files for steelhead and single-end BAM files for Chinook. Two-dimensional SFS and global F ST (weighted) between each population pair were then estimated using realSFS (Korneliussen *et al.* 2014).

To calculate Watterson's theta (Watterson 1975), Tajima's theta (Tajima 1983), and Tajima's D (Tajima 1989), SFS estimated as described above were used as priors (pest) with paired-end BAM files to calculate each statistic for each site (doThetas) which were averaged to obtain a single value for each statistic (Korneliussen *et al.* 2013). The analysis was restricted to 565,000 – 785,000 bp of scaffold79929e for the *GREB1L* region analysis.

The coalescent simulation program ms (Hudson 2002) was used to determine 95% confidence intervals for the theta estimates from 10,000 simulations under a neutral demographic model. The input number of chromosomes was equal to the number of individuals used to calculate the theta statistics. For genome-wide confidence intervals, 100 independent loci and an input theta of 1, which is the approximate theta of a single RAD tag, were used. For the *GREB1L* region confidence intervals, a single locus and the empirical theta estimates were used. The significance of the empirical Tajima's D value was evaluated by generating a Tajima's D distribution from 10,000 ms simulations under a neutral demographic model. A single locus and the average between empirical values of Watterson's and Tajima's thetas in the *GREB1L* region were used. A Tajima's D distribution was also generated using the extremes of the theta confidence intervals and the empirical value remained significant.

Allele frequencies were estimated (doMaf 1; Kim *et al.* 2011) for the significant Chinook SNPs in each population that had at least 4 individuals with enough alignments for subsampling. Paired-end BAM files were used with the reference genome assembly as the prespecified major allele (doMajorMinor 4). Since some populations had low samples sizes, all samples were included regardless of alignment number.

### Amplicon analysis

Amplicon sequence data were demultiplexed by requiring perfect barcode and primer matches. Sequences were aligned to the reference genome assembly described above using the BWA-SW algorithm (Li and Durbin 2010) with default parameters and SAMtools was used to sort, filter for proper pairs, and index BAM files (Table S5).

Phylogenetic analysis was performed on samples in which two or more amplicons had at least 20 alignments (Table S1, S5). Genotypes for all sites were called using ANGSD with the SAMtools genotype likelihood model, a uniform prior, and a posterior cutoff of 0.8. The genotype output file was parsed and converted into biallelic consensus sequences with IUPAC nucleotide code denoting heterozygous positions. These consensus sequences were input into fastPHASE (Scheet and Stephens 2006) to produce 1000 output files that each contained two phased haplotype sequences per individual. Default parameters were used except a distinct subpopulation label was specified for each of the five locations and base calls with a posterior of less than 0.8 were converted to Ns. Parsimony trees were then constructed from each fastPHASE output and a consensus tree was called using PHYLIP (Felsenstein 1989).

In the initial phylogentic analysis, one sample from the Eel River that was originally classified as premature migrating clustered in the mature migration clade (Table S1). A PCA analysis specific to the Eel River placed this sample at an intermediate position between mature migrating and premature migrating sample groups. Furthermore, this was the only Eel River sample that was homozygous for a haplotype associated with residency (Pearse *et al.* 2014). Examination of the original sampling information revealed that this fish was much smaller than others and collected upstream from the main premature steelhead holding area (Clemento III 2006), indicating that it was a resident trout as opposed to an anadromous steelhead. Therefore, this sample was removed and the analysis was rerun.

### Scaffold79929e assembly and annotation

Our initial RAD analysis was aligned against a published reference genome assembly (Berthelot *et al.* 2014) and identified highly associated SNPs on three independent scaffolds. Given the state of the assembly, the sizes of the scaffolds with highly associated SNPs, and the positions of the highly associated SNPs on the scaffolds, we hypothesized that these scaffolds might be physically linked despite not being connected in the current assembly. We aligned four large-insert mate-pair libraries to the published assembly to look for linkages and estimate the distance between linked scaffolds (Table S9). A perfect sequence match was required and alignments to regions with high coverage were discarded. The resulting alignments from all libraries strongly supported a linear assembly with a total size of 1,949,089 bp that included the three associated scaffolds as well as four others (Table S10, S9). This assembled scaffold was named scaffold79929e (e for extended), added to the published assembly, and the seven independent scaffolds that composed it were removed to create the modified reference assembly used in this study.

Scaffold79929e was annotated with MAKER (Cantarel *et al.* 2008) using rainbow trout and Atlantic salmon (*Salmo salar*) EST sequences from the NCBI database, the UniProt/Swiss-Prot database for protein homology, a rainbow trout repeat library (Berthelot *et al.* 2014) for masking, AUGUSTUS (human) and SNAP (mamiso) gene predictors, a maximum intron size of 20,000 bp for evidence alignments, and otherwise default parameters.

### Klickitat steelhead analysis

Single-end RAD data from 237 Klickitat River steelhead samples (Hess *et al.* 2016) was aligned to the modified rainbow trout genome as described above. SAMtools (Li *et al.* 2009) was used to remove unaligned reads, sort, index, and randomly subsample BAM files to 500,000 reads to reduce the effect of PCR duplicates (Andrews *et al.* 2014). All subsequent analyses were performed on subsampled BAM files using ANGSD (Korneliussen *et al.* 2014).

Association mapping was performed using the score test (doAsso 2) with migration date at Lyle Falls (May 1^st^ set to day 1; Hess *et al.* 2016) as a quantitative phenotype (yQuant). The positions of each sample along the first 9 PCs were used as covariates to correct for population stratification. The PCA used to generate covariates was performed as described above.

Genotype data from the four associated SNPs were used to categorize individuals as homozygous for the mature migration allele, heterozygous, or homozygous premature. Genotypes were called (doGeno 4) with a uniform prior (doPost 2) and a posterior probability cutoff of 0.8 (postCuttoff 0.8). 751 out of 948 genotypes passed this cutoff. Two SNPs were flanking sites on the same RAD tag, had near perfect consistency between genotype calls, and were treated as a single genotype for categorization. For an individual to be categorized as homozygous or heterozygous, all called genotypes were required to be in agreement and at least two out of the three genotypes must have been called. A total of 158 samples passed these requirements whereas 51 failed because less than two genotypes were called and 28 failed due to disagreement between called genotypes. Migration date means were calculated with May 1^st^ set to day 1 (Hess *et al.* 2016) and confidence intervals were calculated by bootstrapping with 1000 replicates. The significance of differences in mean migration date between genotype categories was evaluated with Welch’s *t* -test.

### Data availability

The raw sequence data from this study are available at the NCBI Sequence Read Archive with identifier XXXXXXXXX. Scripts to execute the software used for data analysis are available by request to the corresponding author.

## Acknowledgments

The authors thank Chong Liu, Josh Saxon, Peter Tronquet, National Marine Fisheries Service, Oregon Department of Fish and Wildlife, and California Department of Fish and Wildlife for assistance with sample acquisition as well as Fred Allendorf, Eric Anderson, Jamie Ashander, Graham Coop, and Robin Waples for valuable comments on an earlier version of the manuscript. M.R.M. thanks Chris Doe, Yniv Palti, and Gary Thorgaard for invaluable support and encouragement. This work used the Vincent J. Coates Genomics Sequencing Laboratory at the University of California at Berkeley, supported by NIH S10 Instrumentation grants S10RR029668 and S10RR027303.

---

**Figure S1.**
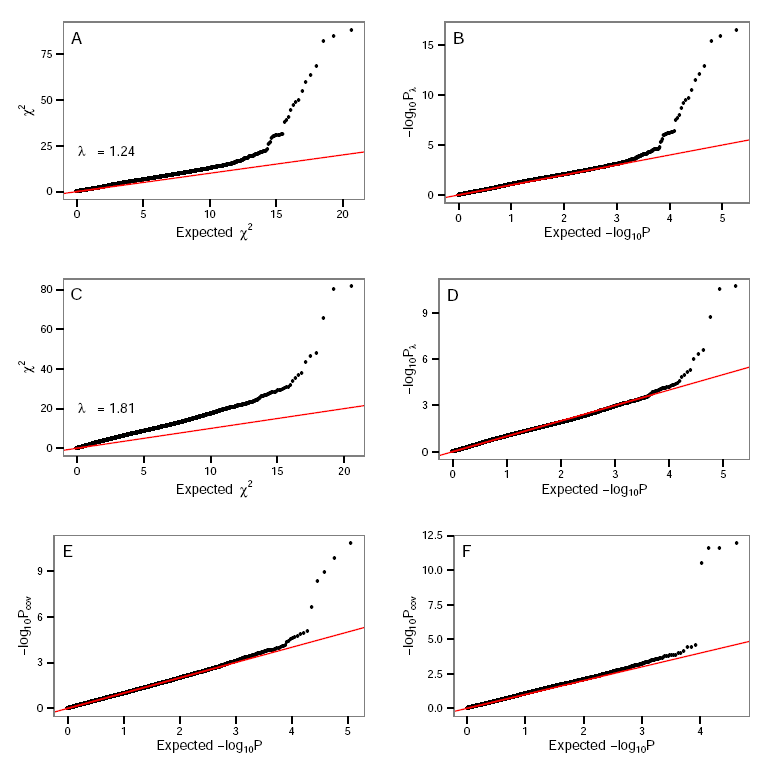
Observed versus expected association statistics in steelhead and Chinook. Chi-squared values and lambda-corrected p-values for North Umpqua (A and B) and Eel (C and D) steelhead. Covariate corrected p-values for all Chinook (E) and Klickitat steelhead (F).

**Figure S2.**
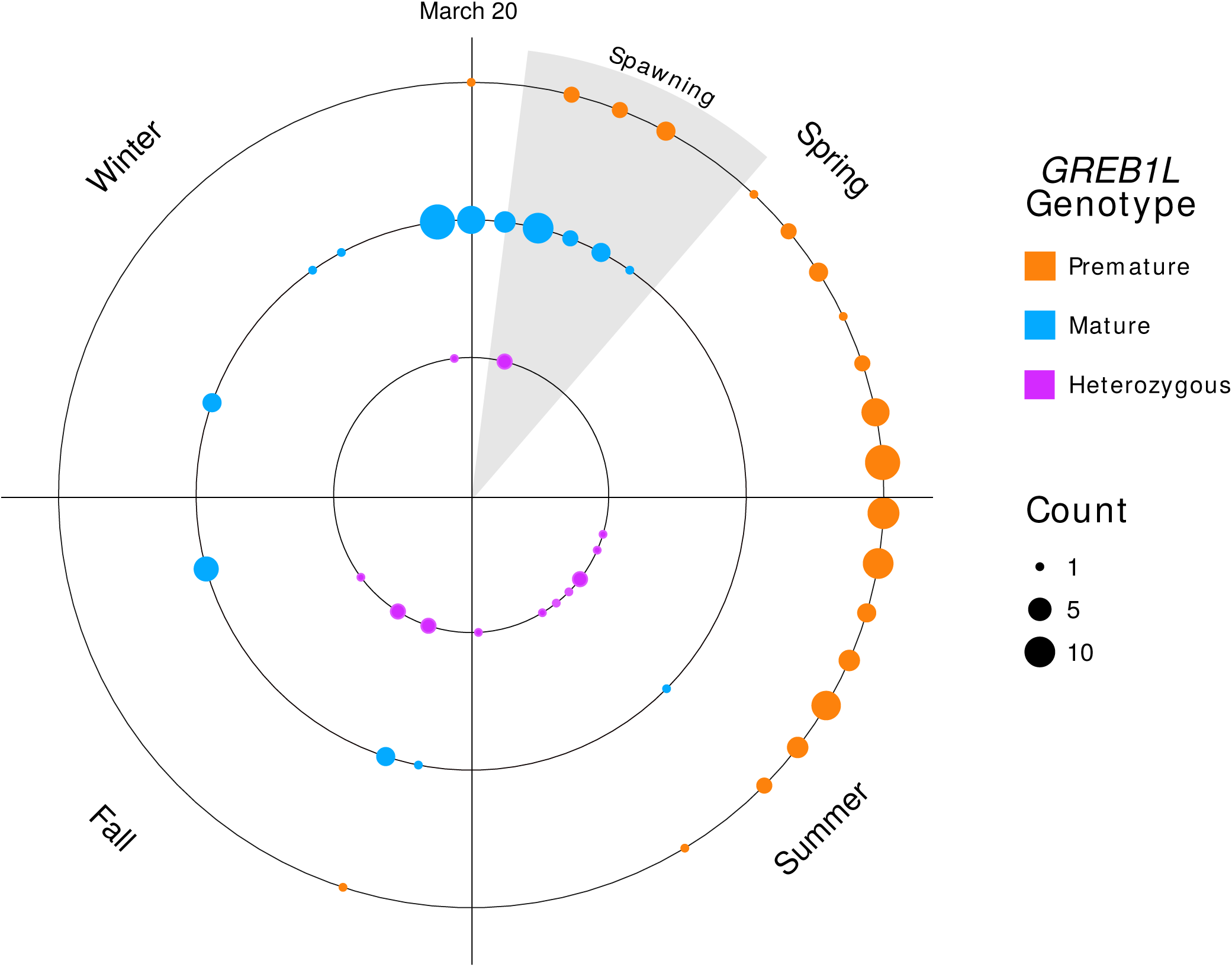
Migration date distribution of Klickitat River steelhead at Lyle Falls with weekly binning. Migration date at specific sites is used as a quantitative proxy for the premature migration phenotype because more direct measures (e.g. gonadal maturation and body fat content at freshwater entry) would be difficult to obtain and require lethal sampling. Individuals categorized as homozygous for the premature allele start migrating at the beginning of Spring and continue through mid-Summer. Premature migrating individuals spawn the year following their migration. Homozygous mature individuals typically migrate in Winter and early-Spring and will spawn the same year they migrate. Most heterozygotes migrate in late-Summer and early-Fall soon after the peak of premature migration. A small number of heterozygotes migrate in Spring, but their gonadal maturation and spawning year is unclear.

